# The Metabolites NADP^+^ and NADPH are the Targets of the Circadian Protein Nocturnin (Curled)

**DOI:** 10.1101/534560

**Authors:** Michael A. Estrella, Jin Du, Li Chen, Sneha Rath, Eliza Prangley, Alisha Chitrakar, Tsutomu Aoki, Paul Schedl, Joshua Rabinowitz, Alexei Korennykh

## Abstract

Nocturnin (NOCT) is a rhythmically expressed protein that regulates metabolism under the control of circadian clock. It has been proposed that NOCT deadenylates and regulates metabolic enzyme mRNAs. However, in contrast to other deadenylases, purified NOCT lacks the deadenylase activity. To identify the substrate of NOCT, we conducted a mass spectrometry screen and report that NOCT specifically and directly converts the dinucleotide NADP^+^ into NAD^+^ and NADPH into NADH. Further, we demonstrate that the *Drosophila* NOCT ortholog, Curled, has the same enzymatic activity. We obtained the 2.7 Å crystal structure of the human NOCT**•**NADPH complex, which revealed that NOCT recognizes the chemically unique ribose-phosphate backbone of the metabolite, placing the 2’-terminal phosphate productively for removal. We provide evidence for NOCT targeting to mitochondria and propose that NADP(H) regulation, which takes place at least in part in mitochondria, establishes the molecular link between circadian clock and metabolism.

## Introduction

The circadian clock adjusts metabolism and behavior of living organisms according to day and night periodicity^1,2^. A screen for circadian genes using *Xenopus laevis* retinas identified the ~50 kDa protein Nocturnin (NOCT), which exhibited a rhythmic expression that peaks at night (16 hours Zeitgeber time, ZT16)^3^. The emergence of sequenced genomes revealed that NOCT is conserved from flies to humans^4–6^. It has been determined that the first description of the NOCT gene happened more than a hundred years ago during studies of *Drosophila melanogaster* (fruit fly) by Thomas Hunt Morgan. The fruit fly NOCT ortholog was called Curled (*cu*) due to a peculiar upward wing curvature in *cu* flies^4^. The *cu* mutant has become a widely used marker in fruit fly genetics and the curled phenotype has been linked to defects in metabolism^4^. In mice, NOCT mRNA undergoes a high-amplitude regulation by circadian clock, peaking by two orders of magnitude in the liver in the transition to evening (ZT12)^7^. NOCT^−/−^ mice exhibit altered lipid metabolism on a high fat diet and do not develop fatty liver disease (hepatic steatosis) or obesity that occur in WT mice^5^. The NOCT^−/−^ mice have low sensitivity to insulin and glucose, and have altered nitric oxide signaling, indicating a deep integration of this circadian protein in mammalian metabolism^5,8–10^.

NOCT is a member of the Exonuclease/Endonuclease/Phosphatase family of proteins which include PDE12, a mitochondrial deadenylase needed for maturation of mitochondrial tRNAs, and CNOT6L, the mammalian ortholog of the main cytosolic yeast deadenylase CCR4^11,12^. Crude NOCT preparations were reported to deadenylate mRNA *in vitro*, leading to the model that NOCT is a circadian deadenylase acting on mRNAs^3^. Regulation of insulin sensing, nitric oxide signaling and lipid metabolism is therefore attributed to a globally altered stability of metabolic enzyme mRNAs^5,8,10^. As a growing body of literature linked the biological effects of NOCT to mRNA deadenylation, two reports described the lack of deadenylase activity in highly purified NOCT *in vitro*^6,13^. Under the same experimental conditions, PDE12 and CNOT6L were active and readily degraded poly-A RNA^6,13^.

To understand the lack of poly-A RNA cleavage by human NOCT, both groups determined the crystal structure of its catalytic domain. The structural studies could not explain the paradoxical inactivity of NOCT, revealing the same fold and the catalytic center as in PDE12 and CNOT6L^6,13^. It has been proposed that NOCT may require a protein partner perhaps for recruitment to mRNAs. Alternatively, it has been postulated that NOCT may have a substrate distinct from poly-A RNA. Here we confirm the second model and show direct cleavage of two chemically related metabolites by human NOCT and by its fruit fly ortholog, Curled.

## Results

### NADP^+^ and NADPH are the direct substrates of NOCT

Based on the absence of mRNA deadenylase activity in purified NOCT, and its known role in metabolism, we hypothesized that NOCT could cleave a metabolite. A previously reported candidate-based approach was unable to find the NOCT substrate^13^, therefore we used an unbiased screen based on metabolomics mass spectrometry. We extracted metabolites from a freshly obtained bovine liver, and independently, from human embryonic kidney HEK293T cells (Fig. 1A). We treated the metabolite extracts with the catalytic domain of recombinant human NOCT (Fig. S1A). NOCT E195A point mutant disrupting magnesium coordination^6^ and WT human deadenylase CNOT6L (residues 158-555) were used as negative controls. Liquid chromatography-mass spectrometry analysis (LC-MS) of the samples revealed that two related metabolites, NADP^+^ and NADPH were selectively depleted by WT NOCT (Fig. 1B; Data Set S1). Their depletion coincided with enrichment in NAD^+^ and NADH (Fig. S1B). These results were reproduced using pure commercially available NADP^+^ and NADPH: the addition of WT NOCT catalytic domain, but not the E195A mutant converted NADPH into NADH and NADP^+^ into NAD^+^ (Fig. 1C). Thus, NOCT uses NADP^+^ and NADPH as substrates and removes the terminal 2’-phosphate (Fig. 1D). To facilitate further analyses of the NADP(H) cleavage reaction, we substituted LC-MS with ion exchange chromatography (IEC). Individual nicotinamide metabolites could be resolved by this technique and quantified by UV absorbance at 260 nm. The IEC assay showed a time-dependent quantitative conversion of both NADP^+^ and NADPH into NAD^+^ and NADH, respectively by full-length (FL) NOCT (Fig. 2A). The activity was uniquely present in NOCT, both full-length and catalytic domain, and absent in the control reactions with human deadenylases CNOT6L and PDE12 (Fig. 2B).

**Figure 1.**
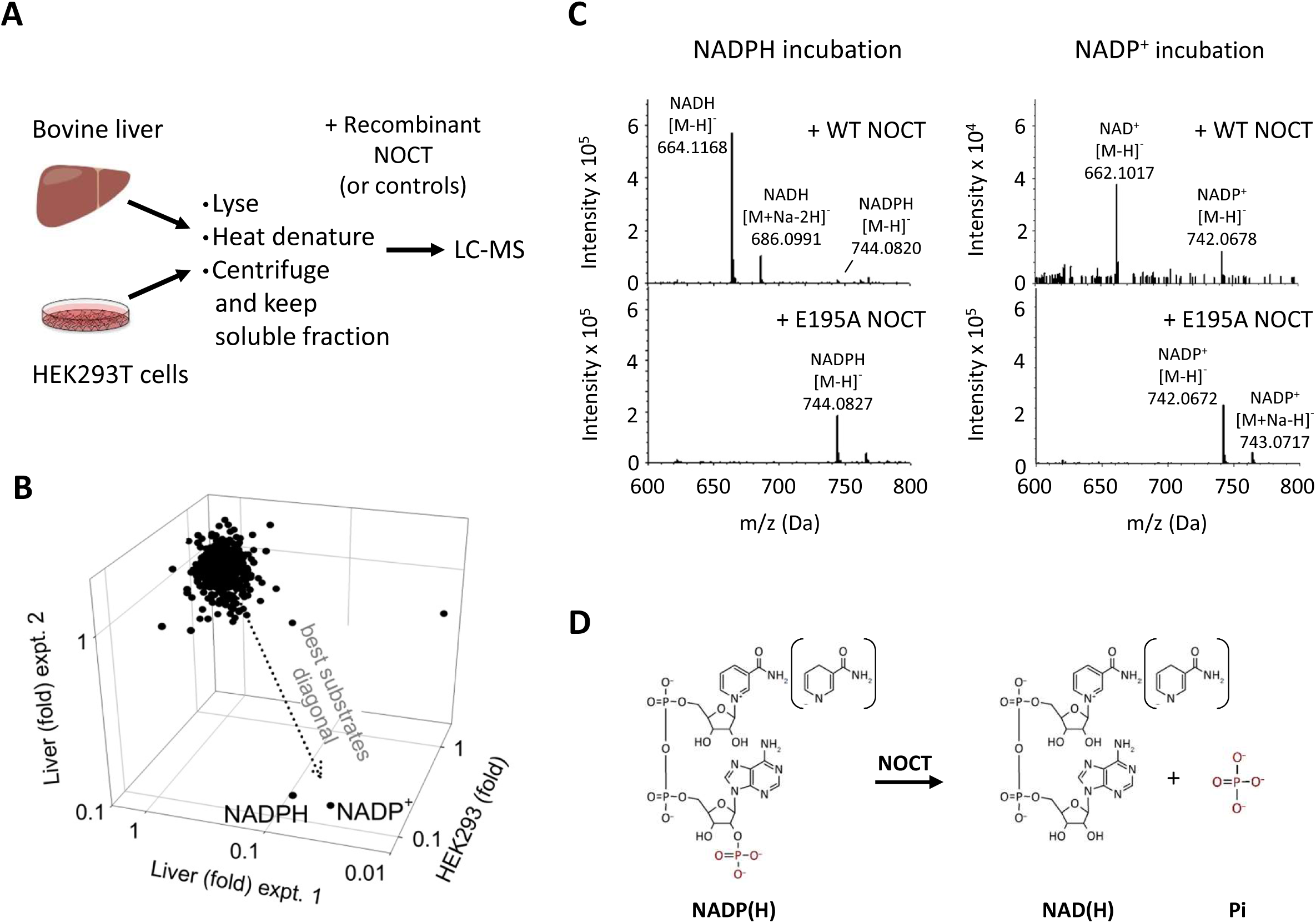
Metabolite profiling to identify NOCT substrate. **A)** Metabolite extraction and treatment outline. Full-length human NOCT and the catalytic domain of NOCT (residues 122-431) was compared against full-length human NOCT E195A, NOCT (122-431) E195A mutant, or the catalytic domain of CNOT6L (158-555). Liver experiment 1 and HEK293 experiment used the catalytic domain of NOCT. Liver experiment 2 used GST-tagged full-length NOCT immobilized on glutathione beads. **B)** Metabolite depletion in three independent experiments (Data Set S1). Fold is defined as the ratio of LC-MS intensities for samples with WT NOCT vs average for samples with NOCT E195A and CNOT6L. **C)** LC-MS/MS analysis of purified NADP^+^ and NADPH following incubation with WT or E195A human NOCT (122-431; 10 μM, 1 hour at r.t.). **D**) The identified reactions catalyzed by human NOCT.

Using conditions of kinetic competition that are established by equimolar mixture of NADP^+^ and NADPH, we determined that NOCT has a small preference for NADPH (Fig. S2A). A titration experiment revealed that the catalytic domain of NOCT cleaves NADPH with K_m_ of approximately 170-300 μM and the first-order catalytic constant k_cat_ = 3.6 s^−1^ for the catalytic domain (Fig. 2C) and 34 s^−1^ for the full-length protein (Fig. S2C). The K_m_ is comparable to that of other enzymes involved in NADP(H) metabolism. The K_m_ of NOCT is within 2-3-fold from that of the recently identified cytosolic NADP(H) phosphatase MESH1 (K_m_ = 0.11 mM)^14^. Further, the NOCT K_m_ is about 5-10-fold better than that for the kinases NADK and NADK2 that synthesize NADPH from NADH (1.7-3.6 mM)^15^. The phosphatase activity observed in NOCT purified from a bacterial expression system was reproduced with overexpressed full-length human NOCT purified from human cells (Fig. 2D-E; Fig. S2B).

**Figure 2.**
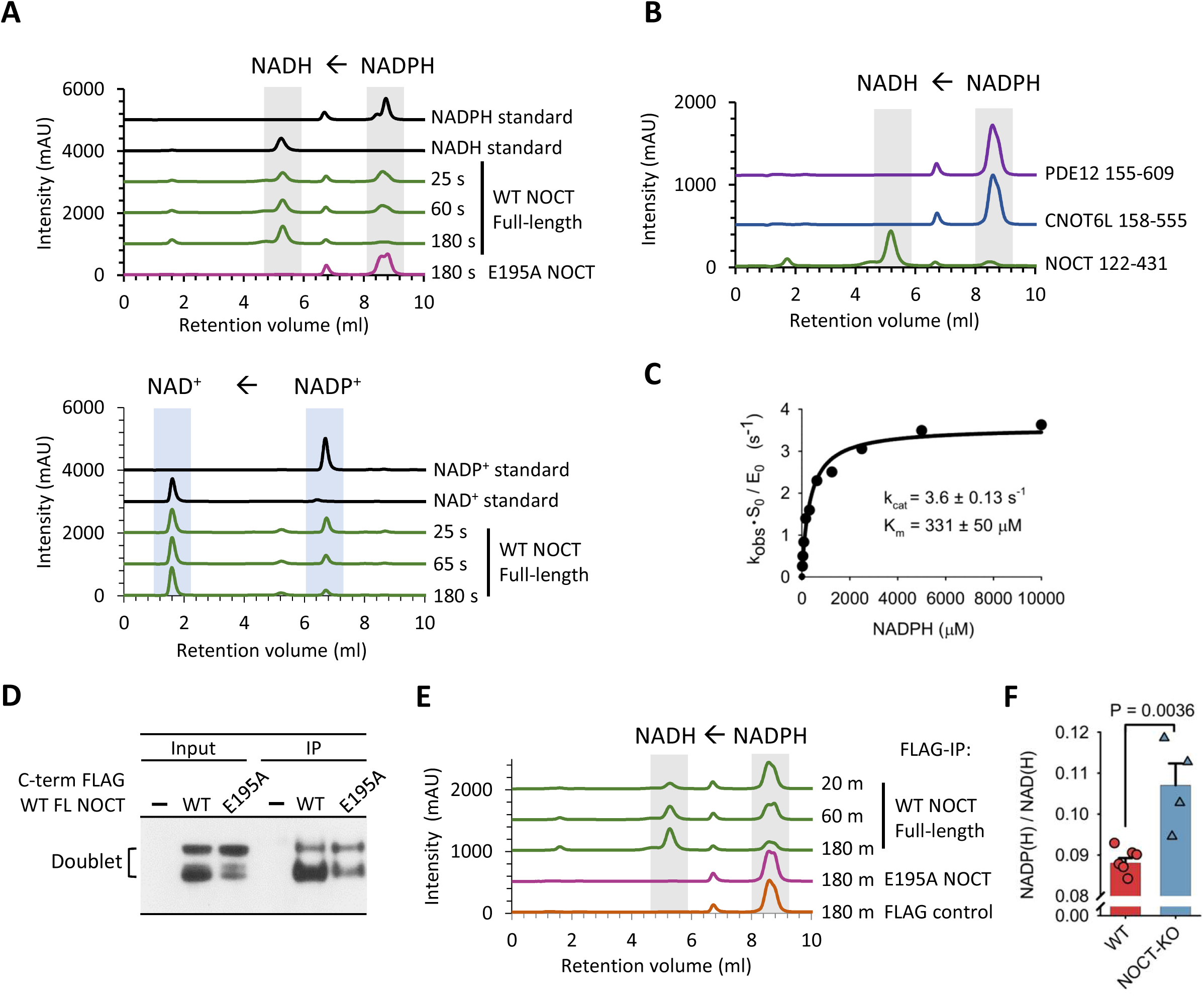
NADP(H) cleavage characterization. **A**) Time course cleavage of pure NADPH (1 mM) and NADP^+^ (1 mM) by purified human full-length NOCT (0.5 μM) at 22 °C, monitored by ion-exchange chromatography (IEC). **B**) 60-minute NADPH incubation with the catalytic domains of PDE12, CNOT6L and NOCT as in (A). **C**) Measurement of kinetic parameters for the catalytic domain of human NOCT (4.8 μM). **D**) Western blot analysis of full-length human FLAG-NOCT (C-terminal tag) expressed in A549 cells for 18 hrs. **E**) NADPH (1 mM) cleavage by FLAG-NOCT purified from human cells. **F**) Nicotinamide metabolite composition in WT and NOCT^−/−^ cells (Data Set S2). Six replicates from three WT samples are compared against four replicates from two independently generated NOCT^−/−^ clones. Error bars are S.E.

To probe regulation of cellular NADP(H) by NOCT, we generated NOCT-KO A549 human cell lines and purified metabolites to determine the NADP(H)/NAD(H) ratios. Faithful capture and quantitation of nicotinamide metabolites in subcellular organelles remains challenging, therefore we carried out whole-cell LC-MS. Analysis of the total cellular metabolites showed a modest, but significant whole-cell increase for NADP^+^ and NADPH in NOCT-KO cells, compared to the 2’-dephosphorylated NAD forms (Fig. 2F; Data Set S2). This experiment may be underestimating the magnitude of 2’-phospho NAD changes inside individual organelles.

### NOCT is targeted to mitochondria

During expression of NOCT in human cells, we noticed that the protein migrated as two discrete bands, which is indicative of cellular processing (Fig 2D). To understand this observation, we used bioinformatics and found that NOCT has a predicted mitochondrial targeting sequence^16^ (MTS, Fig. 3A). The same analysis can discern the presence of an MTS or lack thereof in known mitochondria-targeted proteins or cytosolic proteins, respectively. To test whether the MTS of human NOCT is functional, we generated the deletion mutant, ΔMTS-NOCT. This variant migrated as one band corresponding to the size of processed NOCT (Fig. 3B). Mitochondrial localization of NOCT was supported by confocal microscopy and independently, by subcellular fractionation of overexpressed protein. NOCT was also found in the nuclear fraction based on fractionation but not strongly observed by microscopy. This can be due to a previously reported perinuclear localization of some of the protein^17^. WT NOCT and the active site mutant E195A localized to mitochondria (microscopy) or mitochondria and the nucleus (fractionation), whereas ΔMTS-NOCT appeared in the cytosol (Fig. 3C-D; S3A-B).

**Figure 3.**
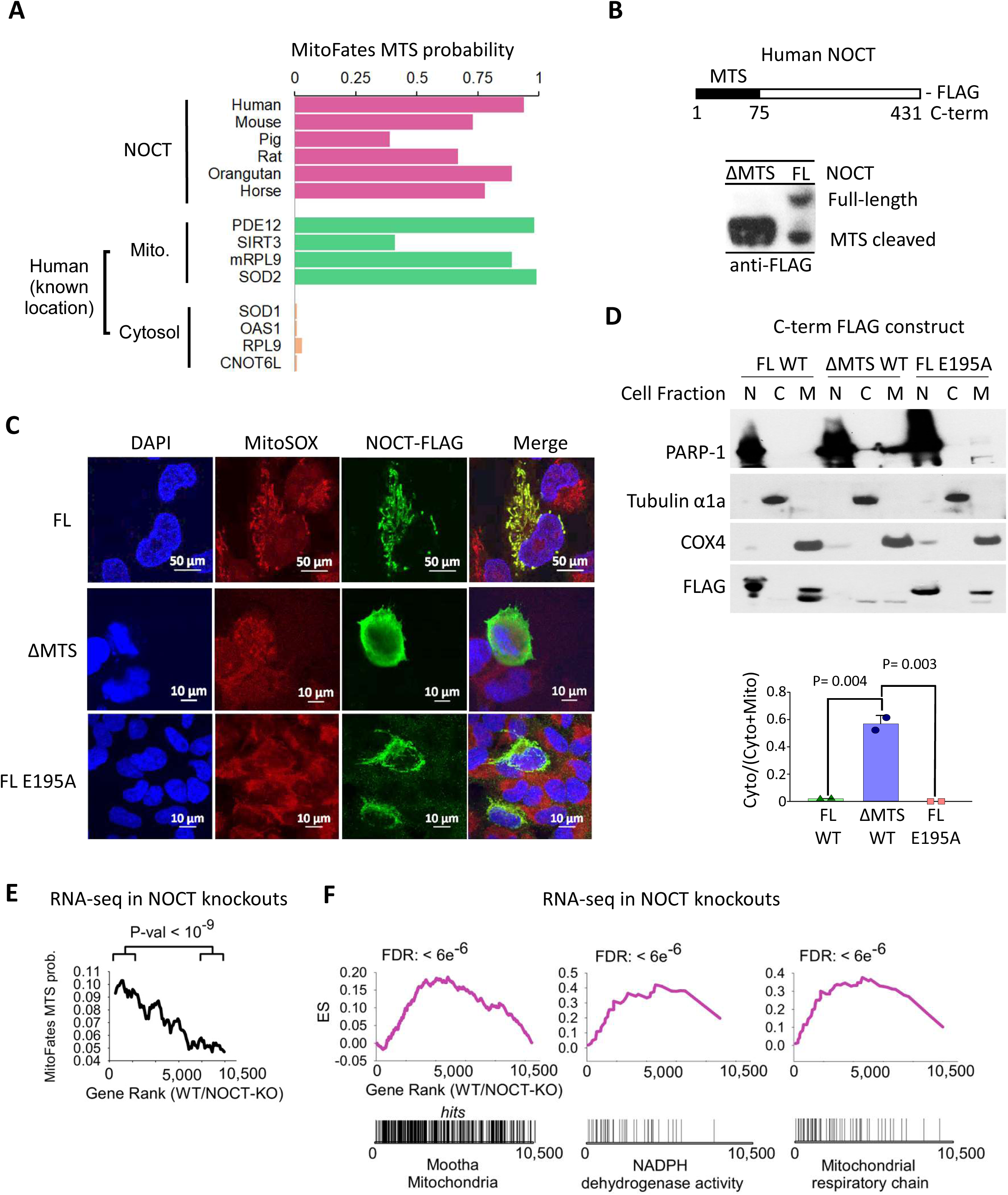
Evidence for NOCT mitochondrial localization. **A**) MitoFates analysis of NOCT and reference human proteins. **B**) Western blot for FLAG-NOCT, full-length vs ΔMTS. **C**) Confocal microscopy of FLAG-NOCT, WT, ΔMTS and E195A active site mutant. DAPI stains for nuclei, MitoSOX stains the mitochondria. **D**) Subcellular fractionation and western blot analysis of NOCT variants in (C). Error bars show S.E. from the data in Fig. 3D and S3B. PARP-1, Tubulin and COX4 mark the nucleus, the cytosol and the mitochondria, respectively. **E**) Running average for MitoFates scores for ~10,500 mRNAs ordered according to gene expression levels in WT vs NOCT^−/−^ cells, measured by RNA-seq (data are provided in Data Set S3). P-values for top 2,000 vs bottom 2,000 mRNAs. RNA-seq data represent four RNA-seq experiments: two in WT cells and two in independently derived NOCT-KO clones. **F**) GSEA enrichment results for the genes with ≥ 50 reads, ranked according to WT/NOCT^−/−^ expression as in (E) (Data Set S3).

To probe the cellular effects of NOCT, we analyzed gene expression differences in WT vs NOCT-KO human cells by RNA-seq (Data Set S3). Compared to NOCT-KO cells, WT cells had increased levels of mRNAs encoding proteins with high-confidence mitochondrial localization based on MTS motifs^16^ and MitoCarta 2 scores^18^ (Fig. 3E; S3C). In agreement with these data, multiple hypotheses testing analysis in GSEA^19^ found gene set enrichments with false discovery rates (FDR) < 0.001 for mRNAs encoding proteins involved in mitochondrial organization and mitochondrial processes (Figs. 3F and S3D). Considering that NOCT cleaves NADP(H) rather than mRNAs, the upregulation of multiple mRNAs in the presence of NOCT is no longer surprising and can be explained by a response to NADP(H) modulation. Together, these RNA-seq results and the presence of an MTS in NOCT point to an important mitochondrial function of NOCT.

### Structure of the complex between human NOCT and NADPH

To understand precisely how NOCT recognizes NADPH, we obtained co-crystals by using calcium instead of magnesium to inhibit the activity of WT NOCT (Fig. S4A) and to stabilize the NOCT**•**NADPH complex. We determined the 2.7 Å crystal structure of the NOCT**•**NADPH complex (Table S1; Fig. S4B) and found the metabolite was bound in a deep pocket comprised of residues that are conserved in the NOCT family^6^ (Fig. 4A-B; Fig. S4C-D). NOCT forms few contacts with the nucleobases. The nicotinamide group does not pack strongly against any NOCT residues and has a weak electron density. The adenine is sandwiched between two arginines (R290 and R367) via cation-pi interactions. The density for the adenine is relatively weak, indicating that these interactions are dynamic (Fig. 4C; S4D). In contrast, the ribose-5’-PP-5’-ribose-2’-P backbone of NADPH forms multiple stable contacts and has strong density. The recognition of the sugar-phosphate backbone appears to be incompatible with other cellular ligands, including ATP, RNAs, mRNA 5’-cap (which has a ribose-5’-PPP-5’-ribose linkage, and 3’-P rather than 2’-P at the scissile position), and NAD^+^-capped mRNAs^20^, which have 3’-P rather than 2’-P at the scissile position.

**Figure 4.**
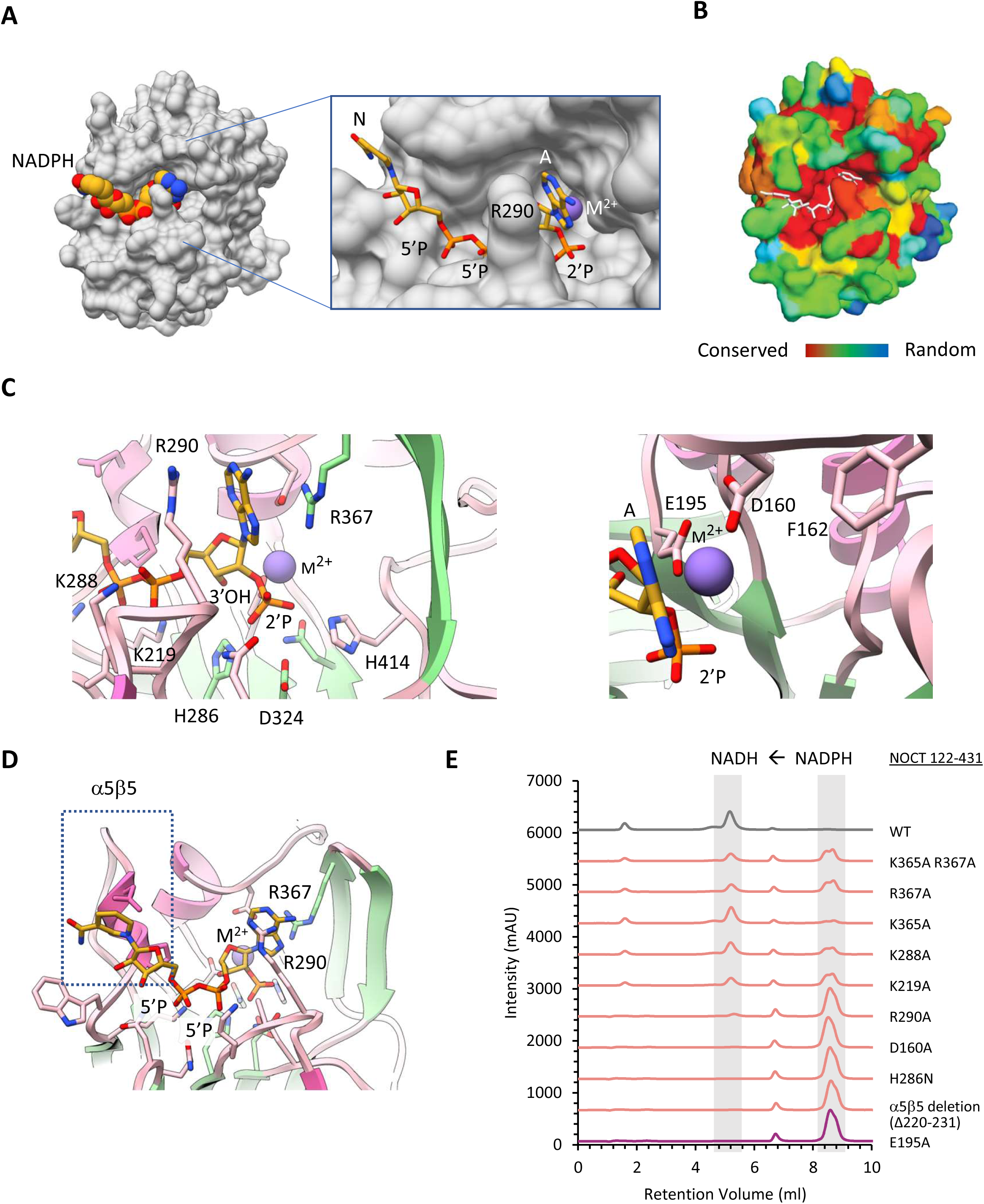
Structure of human NOCT•NADPH•Ca^2+^ complex. **A**) Surface representation of the complex with NADPH bound to the active site pocket. **B**) NOCT conservation from 351 non-degenerate sequences mapped onto the NOCT structure. **C)** NADPH position and the divalent metal ion coordination in the catalytic center. **D**) NADPH position relative to the α5β5 motif. **E**) NADPH cleavage by NOCT mutants designed to disrupt the enzymatic activity. NOCT catalytic domain (500 nM) and NADPH (0.5 mM) were used. Reactions were conducted at 22 °C for 60 minutes.

The 2’-P of NADPH is placed into a chemically ideal position for dephosphorylation (Fig 4C). The chemical step for the ribose-phosphate hydrolysis should be the same as in the related mRNA deadenylases PDE12 and CNOT6L, and the DNA repair enzyme TDP2 due to superposition of their magnesium-coordinating catalytic centers^6,12,21^. For TDP2, which has been crystallized with the DNA, the scissile 5’-phosphate and the catalytic Mg^2+^ ion were positioned with confidence in the active site aided by the size of the DNA substrate. Superimposition of our NOCT**•**Ca^2+^**•**NADPH structure with this TDP2**•**Mg^2+^**•**DNA complex shows identical metal ion and phosphate placements, indicates that the phosphatase activity of NOCT and the phosphodiesterase activity of TDP2 indeed involve the same chemistry (Fig. S4E-F). In NOCT, the phosphate coordinates with the catalytic metal ion, which is positioned to stabilize the leaving group (ribose oxygen). The 3’-hydroxyl is buried inside the active site and away from the metal ion. The orientation of the ribose near the catalytic metal ion places the 2’ ribose position rather than the 3’ ribose position for chemical attack. This can explain the inability of NOCT to cleave RNA. The catalytic water molecule that serves as the incoming nucleophile must be activated by ASP324 (Fig. 4D), which is the only residue with pK_a_ near neutrality next to the nucleophile position. The mode of NADPH recognition by NOCT is unique and bears little resemblance with the only other professional NADP(H) phosphatase described to date, MESH1 (PDB ID 5VXA)^14^. In MESH1 nucleobases rather than the sugar-phosphate backbone form the majority of protein-substrate contacts, whereas the catalytic metal ion Zn^2+^ is placed to activate the nucleophilic water molecule rather than the leaving 2’-O group of the ribose.

Bioinformatics analysis has shown that the helix α5 linked to the strand β5 (α5β5 motif) is a conserved structural feature of NOCT that is not conserved in homologous mRNA deadenylases^6^ (Fig. 4D). In the NOCT**•**NADPH complex, the α5β5 motif packs closely with the nicotinamide-bearing sugar. Superimposing structures illustrates that neither CNOT6L nor PDE12 has the α5β5 motif in the position for binding NADPH (Fig. S4G). Additionally, the mRNA deadenylases contain a tryptophan instead of an alanine present in NOCT (Fig. S4H), which creates a steric clash with the diphosphate-sugar backbone of NADPH and further distinguishes NOCT from mRNA deadenylases. We mutated a number of NOCT active site residues proximal to NADPH, as well as deleted the α5β5 motif and determined that all of these mutations are detrimental to the NADPH 2’-dephosphorylation activity (Fig. 4E).

### *Drosophila Melanogaster* Curled Cleaves NADP^+^ and NADPH

Conservation analysis shows that the NADPH-binding pocket is conserved between human NOCT and Curled, suggesting that our findings with human NOCT may extend to the fly ortholog (Fig. 5A). To test this prediction, we cloned and purified the full-length *Drosophila melanogaster* protein (Fig. S5A) and tested its RNase and 2’-phosphatase activities. We determined that Curled lacks deadenylase activity when tested using radiolabeled poly-A RNA. Neither WT Curled nor Curled E140A active site mutant cleaved poly-A, whereas the control deadenylases CNOT6L and PDE12 degraded the RNA substrate within minutes (Fig. 5B). In contrast, Curled readily cleaved NADPH (Fig. 5C; Fig. S5B). This catalytic activity was blocked by the active site point mutant E140A, which mimics the E195A mutation in human NOCT. Further extending the similarity with human NOCT, Curled cleaved both NADP^+^ and NADPH, exhibiting a preference for NADPH (Fig. S5C). Full-length Curled shows ~2-fold preference for NADPH (Fig. S5C), which is comparable to the 6-fold preference observed with full-length NOCT (Fig. S2A). For NADP^+^ the specific cleavage activity of Curled and NOCT are the same, whereas for NADPH NOCT is ~4-fold more active.

**Figure 5.**
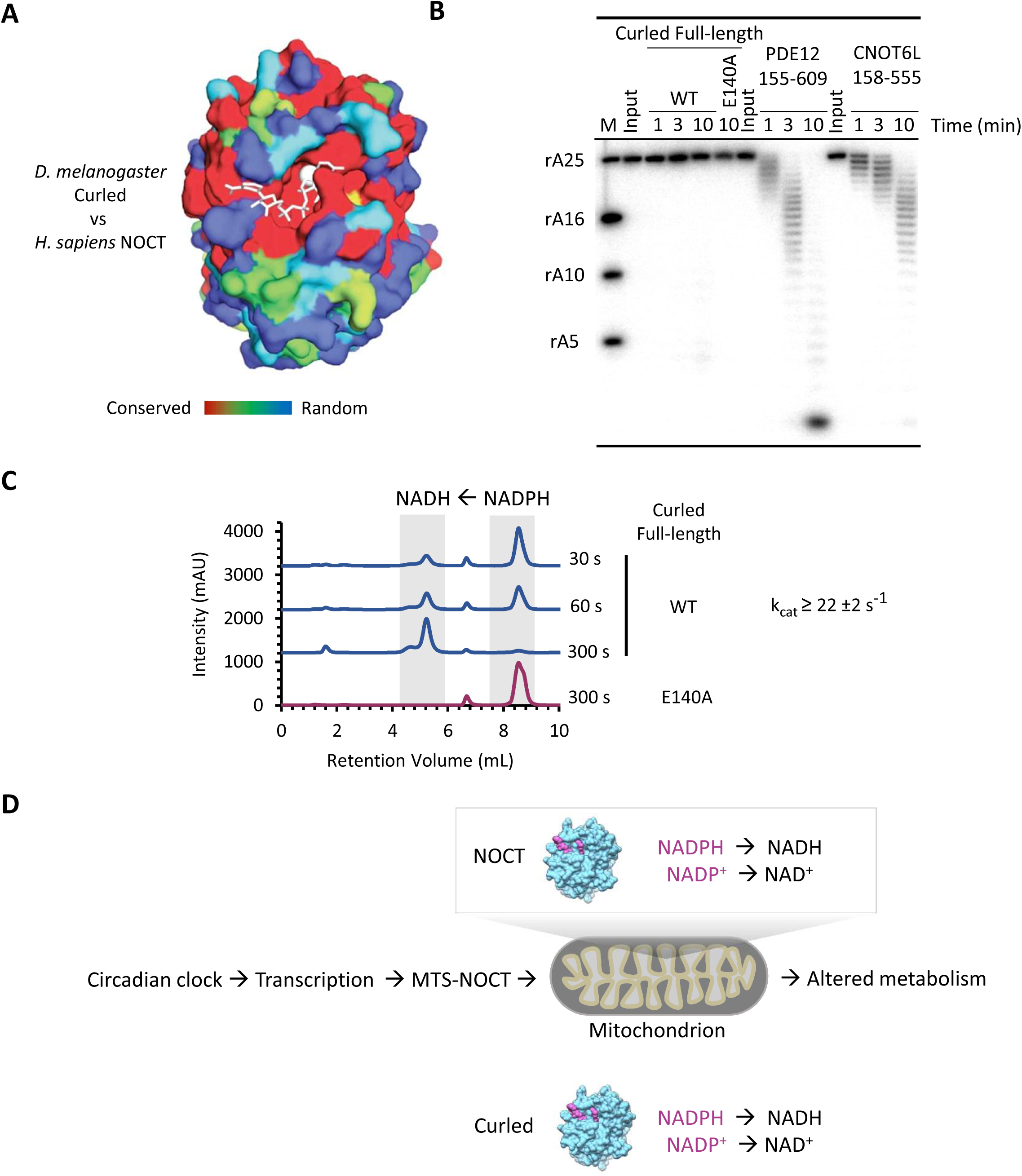
Biochemical characterization of Curled. **A**) Structure of human NOCT colored by sequence conservation between fruit fly and human proteins. **B**) ^32^P-poly-A RNA cleavage assays using *D. melanogaster* Curled, Curled E140A active site mutant, and human deadenylases CNOT6L and PDE12 (2 μM each). Faint bands at 10 minutes in WT Curled arise from a minor contaminating endonuclease. The E140A mutant corresponds to the E195A mutation in human NOCT (Fig. 2A and 4E). **C**) IEC trace for cleavage of NADPH (1 mM) by Curled from *Drosophila Melanogaster* (500 nM). The catalytic constant was calculated as k_obs_**•**S_0_/E_0_ from Figures 5C and S5B. This value represents the lower estimate for k_cat_. **D**) The proposed model for NOCT integration in the circadian system based on the results of our work.

## Discussion

We identified NADP^+^ and NADPH as the substrates of the circadian enzyme, Nocturnin. Although the involvement of NOCT in metabolism has been widely recognized, it had not been expected that this enzyme directly regulates the central cofactors in anabolic and catabolic reactions, NADP^+^ and NADPH (substrates), and NAD^+^ and NADH (products). NOCT does not degrade NADP(H), but converts it into a coupled cofactor NAD(H) essential for energy metabolism^22^. Enzymes that use NADPH co-factors have key roles in mitochondrial oxidation of unsaturated lipids^23^ and regulation of body weight and insulin sensing^24,25^, which also represent the key phenotypes reported for NOCT. Previously, the physiological effects of NOCT have been attributed to metabolic mRNA deadenylation^10^. Our work shifts the focus to the direct target, NADP(H), and thereby establishes a solid ground for re-interpretation and mechanistic understanding of the functions of this enzyme in metabolism and circadian clock control. For example, the sharp peak in NOCT expression when mice wake up (mice are nocturnal and ZT12 corresponds to awakening) must be understood from the point of NADP(H)/NAD(H) regulation. One physiologic function of NOCT could be to maximize available NAD(H) for energy generation in a search for food, using elevated blood sugar that animals have at the time of awakening^26^.

The demonstration that Curled cleaves NADPH reveals the molecular target of the mutant *cu*, which has been used for more than a century in fruit fly genetics. This result bridges a 103-year gap in knowledge and in addition verifies that NADP(H) 2’-dephosphoylation is a conserved activity in the NOCT family. Of note, in 1923, Lenore Ward described a dominant mutation, Curly, which caused upward wing curvature analogous to that seen in Curled. It has been established recently that Curly is not related to Curled and encodes an NADPH oxidase that generates reactive oxygen species (ROS)^27^. Therefore, both genes linked to the curled wing phenotype, Curled and Curly, share the same NADPH substrate, suggesting that they may regulate the same or overlapping pathways controlling wing morphology, and that these activities could depend on ROS signaling.

Human NOCT is encoded by a single isoform with an MTS, which can target the enzyme to mitochondria. At least some of the biological functions of human NOCT therefore involve cleavage of mitochondrial NADP(H) (Fig. 5D). The MTS is present in the NOCT gene of higher animals, but in *Drosophila melanogaster* only two of the six isoforms are predicted to have an MTS. These observations suggest that Curled and perhaps also NOCT may have functions inside and outside of mitochondria. In conclusion, we propose that NOCT/Curled convert NADP^+^ to NAD^+^ and NADPH to NADH to regulate metabolism and in the case of NOCT, to synchronize metabolism with the biological clock.

### Nomenclature considerations

NOCT has Enzyme Commission classification 3.1.13.4: poly(A)-specific 3'-exoribonuclease^28^. Our work shows that NOCT does not position the 3’ hydroxyl productively in the active site and that instead NOCT is an NADP(H) 2’-phosphatase. This activity does not match a present EC classification. At the moment there is only one example of a professional NADP(H) 2’-phosphatase, MESH1, described in a preprint publication^14^. MESH1 does not localize to mitochondria, it has a different structure and a different NADP(H) binding mode than NOCT, and it does not have known associations with the circadian clock or fly phenotypes. We propose that the discovery of two structurally and biologically distinct 2’-phosphatases calls for the creation of an EC entry to describe this enzyme class.

## Materials and methods

### Metabolite extraction and LC-MS

Cow liver (purchased fresh from Cherry Grove Farm, Lawrenceville, NJ, USA) was homogenized using Retsch Cyromill and the homogenate (~0.8 grams) was resuspended in 600 μl sterile water. For HEK293T cells, nine 10 cm dishes containing confluent cells were washed with 2 ml of cold phosphate-buffered saline (PBS). Cells (~0.4 grams) were scraped off and spun down at 100 x g for 5 min in 1.5 ml tube. The supernatant was discarded and the cells were resuspended in 300 μl sterile water. From this point forward liver and HEK293T cells were treated similarly.

The suspensions were sonicated on ice on Branson Sonifier S-450 with 20 pulses (Duty Cycle: 30%, Output: 4). The lysates were heated to 95 °C for 10 min then centrifuged at 22,000 x g for 10 min (4 °C). The cleared lysates were used to set up separate reactions with soluble NOCT, E195A NOCT, CNOT6L, and blank in a total volume of 135 μl (liver) or 90 μl (HEK293T). For bead-immobilized full-length NOCT, 100 μl of bed volume was used per reaction. Each reaction tube was supplemented with 10 μM proteins (f/c) (except blank and bead samples), 0.5 mM MgCl_2_, and incubated for 19 hrs at 22 °C. The obtained samples were analyzed by LC-MS, which detected ~17,000 unique compounds. LC-MS measurements were performed with a quadrupole Orbitrap mass spectrometer (Q Exactive Plus, Thermo Fisher Scientific), operating in negative ion mode, and coupled to hydrophilic interaction chromatography via electrospray ionization. The scan range was *m/z* 73 to 1000.

### Chromatographic NADP(H) 2’-dephosphorylation assays

Analyses of NADP(H) cleavage were carried out at 22 °C using 1000 μM NADP(H) (Roche) and 0.5 μM Nocturnin, unless otherwise noted. Reactions contained 20 mM Tris⋅HCl (pH 8.0), 70 mM NaCl, and 2 mM MgCl_2_. The reactions were stopped at indicated times by quick heating to 95 °C. Samples were injected onto a Mono-Q column on AKTA fast performance liquid chromatography instrument (GE Healthcare Life Sciences) and eluted in 20 mM Tris⋅HCl (pH 8.0) with 0→2 M NaCl gradient. Standards for NAD^+^, NADH, NADP^+^ and NADPH were from Roche. Peak areas were used to quantify metabolite abundances and determine cleavage kinetics.

### NOCT•NADPH co-crystallization

Crystallization drops contained 12 mg/ml of NOCT 122-431, 2 mM CaCl_2_, and 100 mM NADPH (Roche). Calcium was selected instead of magnesium to stabilize the metabolite. Our experiments determined that NOCT is inactive in the presence of Zn^2+^ and Ca^2+^ presumably due imperfect transition state formed due to a slight size mismatch of these ions. Co-crystals were grown using the hanging drop vapor diffusion method by mixing the crystallization complex 1:1 with reservoir solution (100 mM HEPES pH 7.5, 10% (v/v) Isopropanol, 50 mM Sodium Acetate, and 12% (w/v) PEG 4,000). Crystals were cryoprotected using 100 mM HEPES pH 7.5, 10% Isopropanol, 50 mM Sodium Acetate, 12% (w/v) PEG 4,000, 100 mM NADPH, 2 mM CaCl_2_, and 25% (v/v) glycerol, then frozen in liquid nitrogen.

### X-Ray Data Collection and Structure Determination

X-ray diffraction data were collected at the AMX (17-ID-1) beam line at the Brookhaven National Laboratories (NSLS-II). Data were collected at a wavelength of 0.97 Å and processed with XDS and then STARANISO to correct for diffraction anisotropy. Co-crystals contained a single NOCT copy per asymmetric unit and belonged to the tetragonal P4_1_2_1_2 space group. The structure was solved by molecular replacement in PHASER, using the human apo NOCT structure (PDB ID 6MAL) as the search model. Structure of the complex was built in COOT and refined using simulated annealing (5000K, then 2000K for final stages) using PHENIX.

## Acknowledgements

We thank staff at AMX (17-ID-1) and FMX (17-ID-2) beam lines, Dr. Phil Jeffrey for his assistance with data collection and processing, and Prof. Fred Hughson for helping with timely access to the beam lines. We also thank Dr. Gary Laevsky at the Department of Molecular Biology, Princeton University, and *Nikon* Center of Excellence for providing the microscopy equipment, Dr. Wei Wang and staff at Lewis-Sigler Genomics institute RNA-seq facility, and Dr. Tharan Srikumar and Dr. Henry Shwe of the Proteomics and Mass Spectrometry Core. We thank Prof. Mohamed Donia and Dr. Yuki Sugimoto for helpful discussions regarding metabolites, Dr. Girish Deshpande for insights into fly genetics, Dr. Lei Wei from Prof. Alexander Ploss’s Laboratory for help designing CRIPSR KO cells, Guanhua He from Prof. Martin Jonikas’s Laboratory for help with cell homogenization and Xinlei Sheng from Prof. Ileana Cristea’s Laboratory for help with subcellular fractionation.

## Funding

This study was funded by Princeton University, NIH grant 1R01GM110161, Burroughs Wellcome Foundation Grant 1013579 and The Vallee Foundation grants (A.K.), NIH grants 5T32GM007388 and F99 CA212468-01 to S.R., a pre-doctoral fellowship from the China Scholarship Council - Princeton University Joint Funding Program to J.D., NIH grants 1DP1DK113643, R01 CA163591, P30 CA072720 (Metabolomics Shared Resource, Rutgers Cancer Institute of New Jersey), Stand Up To Cancer - Cancer Research UK - Lustgarten Foundation Pancreatic Cancer Dream Team Research Grant (SU2C-AACR-DT-20-16) to J.R, and NIH R35 GM126975 grant to P.S. Work at the AMX (17-ID-1) and FMX (17-ID-2) beam lines is supported by the National Institute of Health, National Institute of General Medical Sciences grant P41GM111244 and by the DOE Office of Biological and Environmental Research grant KP1605010. The National Synchrotron Light Source II at Brookhaven National Laboratory is supported by the DOE Office of Basic Energy Sciences under contract number DE-SC0012704 (KC0401040).

## Competing interests

The authors declare no competing interests.

## Author contributions

M.E. and J.D. generated samples for LC-MS and conducted biochemical studies and crystallization. M.E. built the structure and performed subcellular fractionation. J.D. performed fluorescence microscopy. L.C. designed the metabolite screen, performed LC-MS and analyzed the MS data. S.R. purified and analyzed FLAG-NOCT from human cells. E.P. and A.C. generated A549 knockout cells, and E.P. conducted RNA-seq experiments and RNA-seq data processing. T.A. and P.S. generated fruit fly Curled DNA construct. J.R. proposed the metabolite screen strategy and provided LC-MS technology. A.K. performed GSEA analyses. M.E., J.D., and A.K. wrote the manuscript. A.K. supervised the work.

## Data Availability Statement

RNA-seq data have been submitted to GEO database under ID GSE123477. Structure and the diffraction data have been submitted to PDB database under ID 6NF0. Figures 1B, 2F, 3E-F, S3C-D have raw data provided as Data Sets S1-S3. There are no restrictions to data availability, except that the PDB entry will be released upon publication.

## Supplementary Information

**Figure S1.**
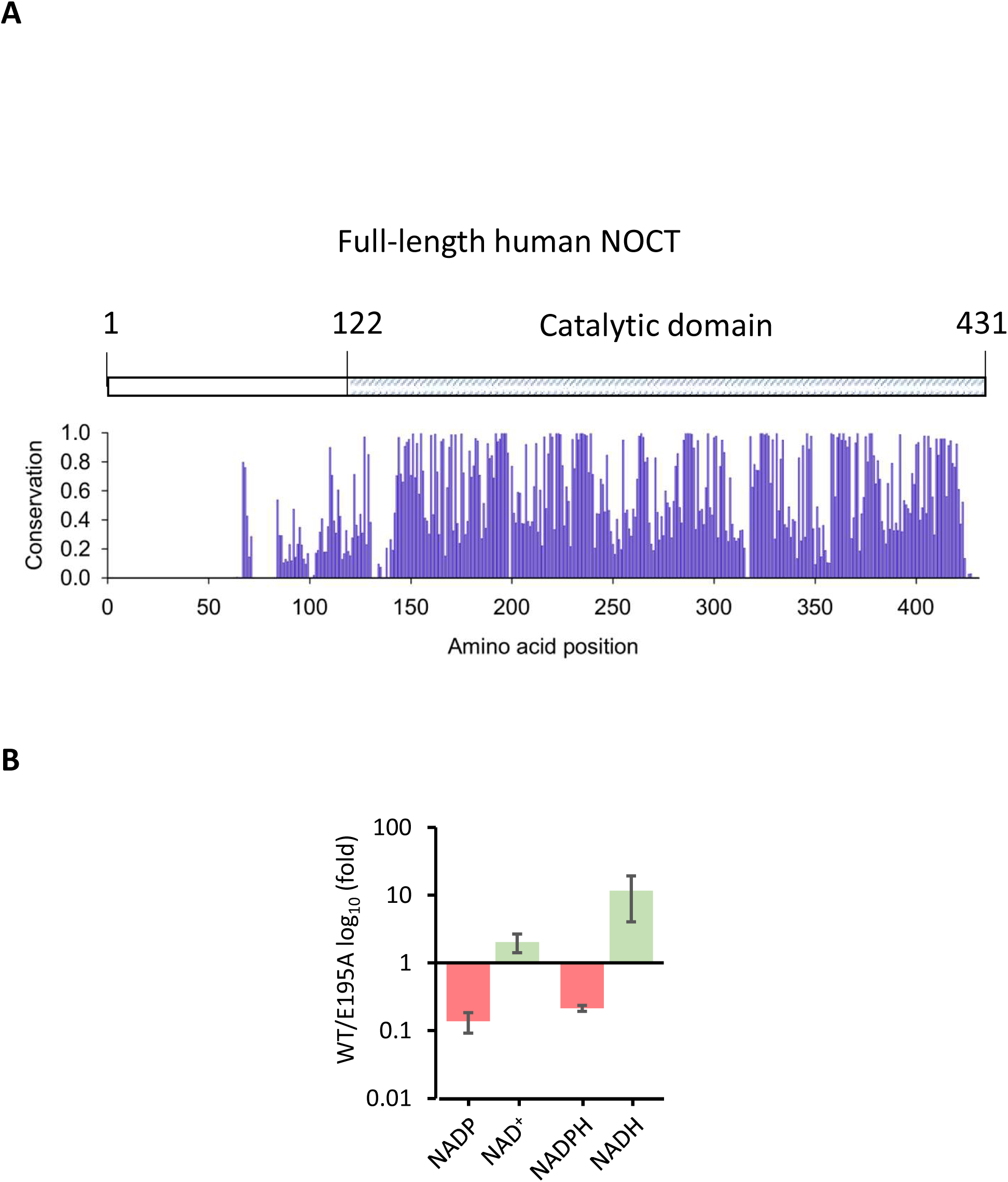
NOCT sequence and metabolite conversion activity. **A)** Sequence alignment for 351 non-redundant NOCT sequences was used to compute the conservation values. Conservation was computed using SEQMOL-Kd (BiochemLabSolutions). **B)** The depletion of NADP(H) in the metabolite screen experiments (Fig. 1A) is accompanied by the expected increase in NAD(H). The basal levels of NAD^+^ considerably exceed those of NADP^+^, which accounts for a smaller magnitude of NAD^+^ increase compared to NADP^+^ loss.

**Figure S2.**
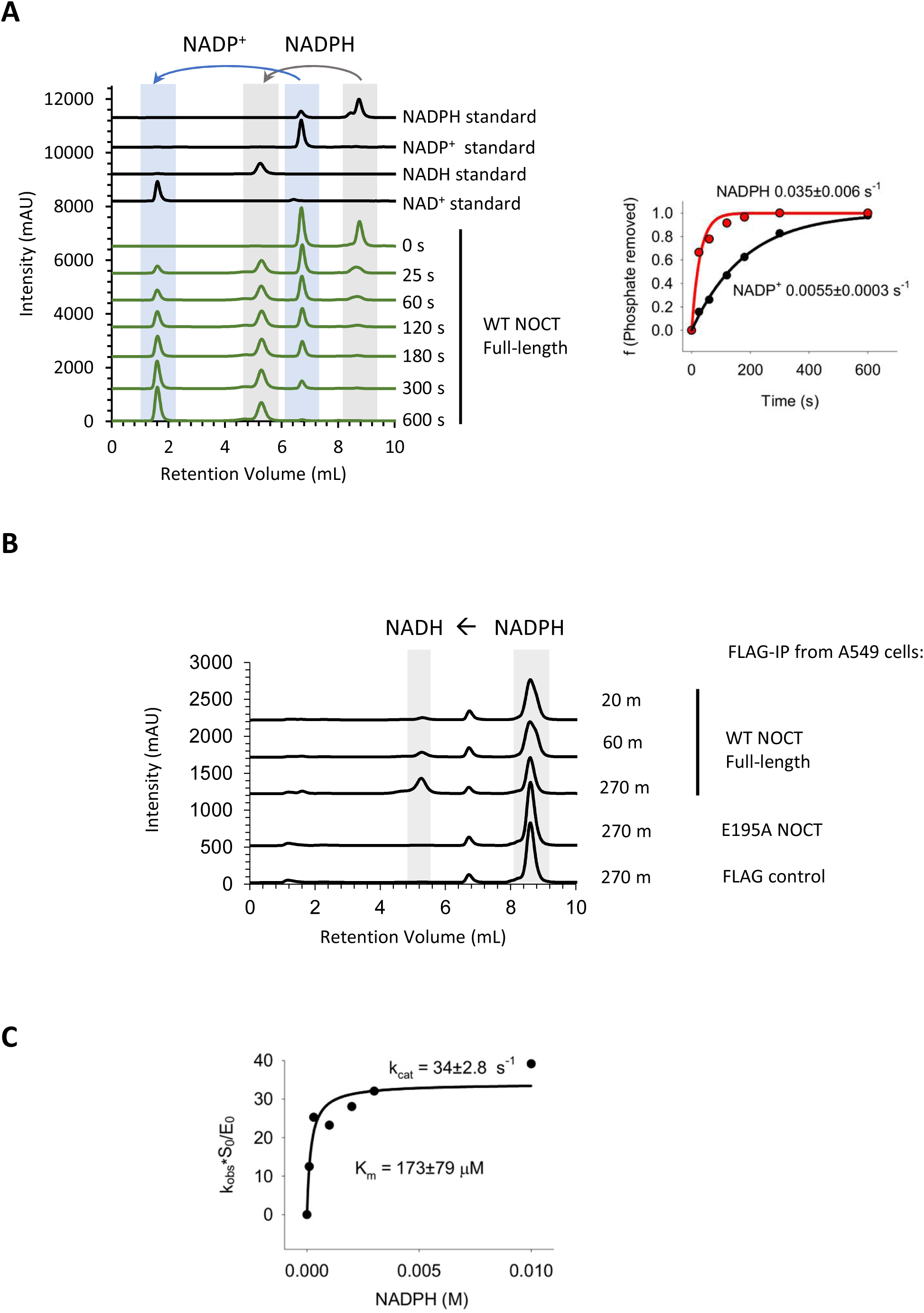
NADP^+^ and NADPH cleavage analyses. **A**) Left: Cleavage of NADP^+^ and NADPH under conditions of kinetic competition. Reactions contained 0.5 μM full-length WT human NOCT and 1 mM of each metabolite. Right: quantitation of the data fitted to single-exponential kinetics. **B**) A biologic replicate of NADPH (1 mM) cleavage by full-length NOCT on beads, which was purified from human cells (related to Fig. 2E). **C**) Kinetic parameters for NADPH cleavage by full length NOCT determined using 0.5 μM NOCT and as in Fig. 2C.

**Figure S3.**
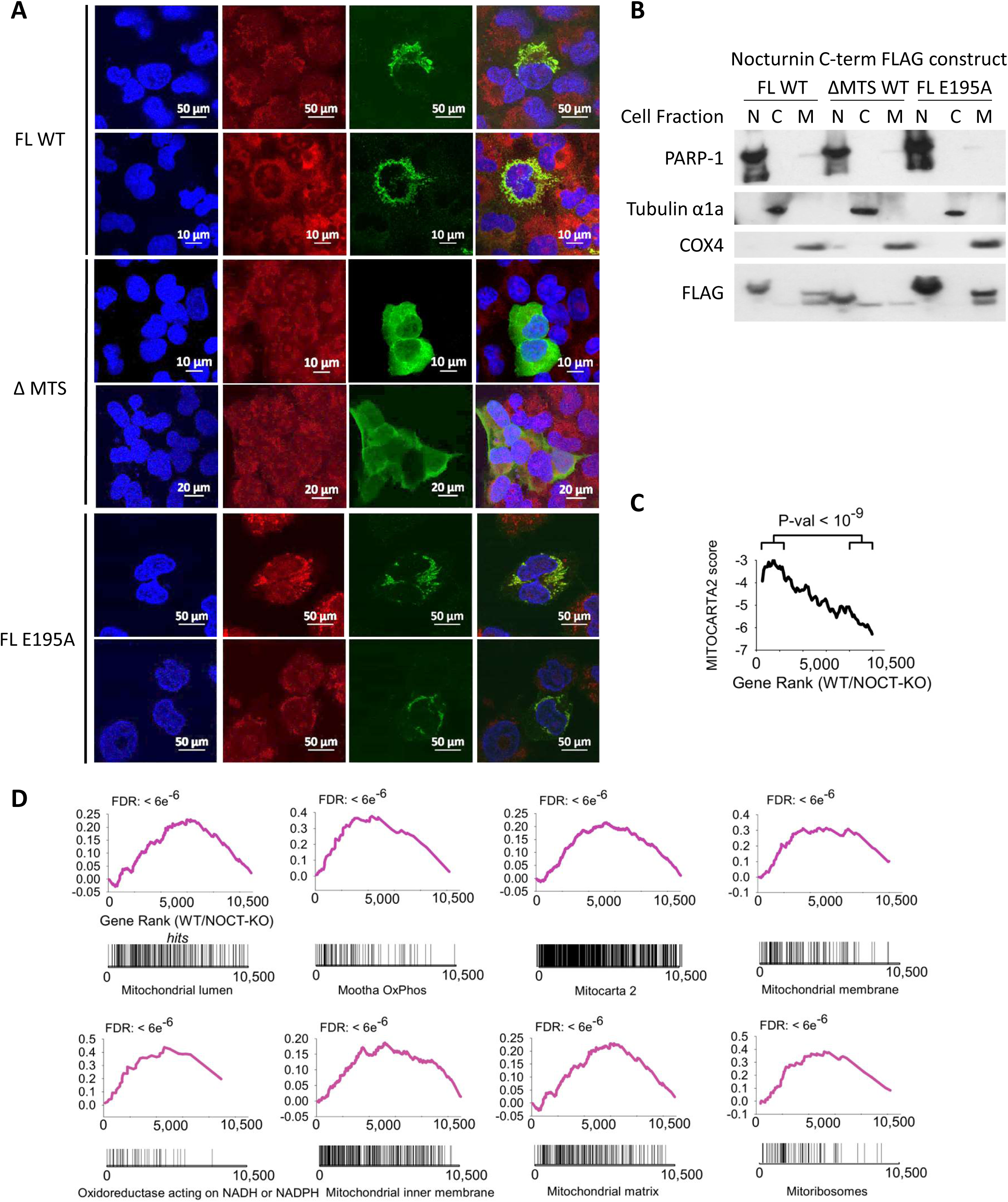
NOCT localization and effect on transcription in human cells. **A**) Confocal microscopy of FLAG-NOCT, WT, ΔMTS and E195A active site mutant related to Fig. 3C. **B**) Subcellular fractionation and western blot analysis of NOCT variants related to Fig. 3D. **C**) Running average of MitoCarta 2 scores for mRNAs upregulated in WT vs NOCT^−/−^ cells, as determined by RNA-seq. The P-values were obtained as in Fig. 3E. RNA-seq data represent four RNA-seq experiments: two in WT cells and two in independently derived NOCT-KO clones. **D**) GSEA enrichment for mitochondria-related terms in WT vs NOCT^−/−^ cells as in Fig. 3F.

**Figure S4.**
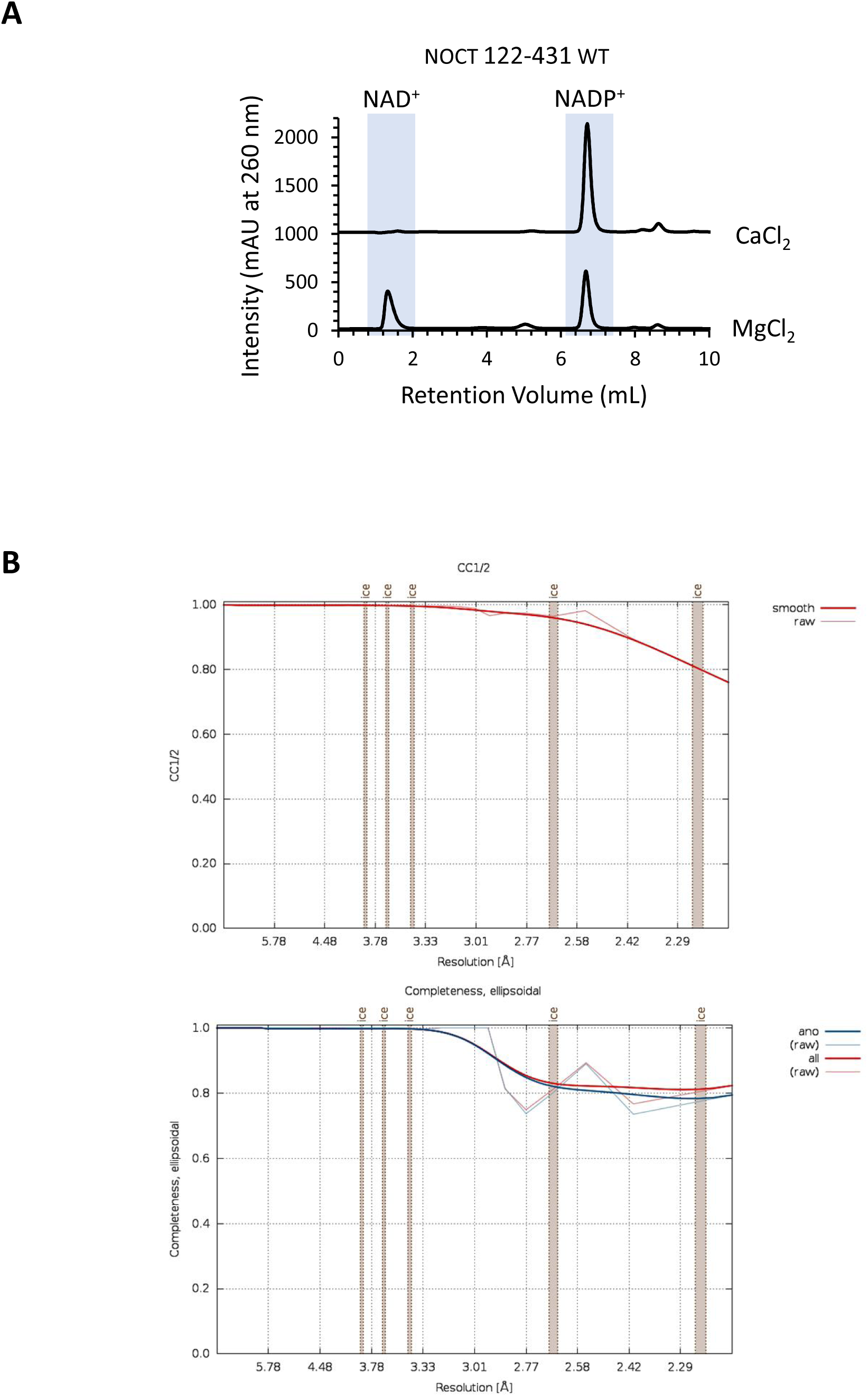
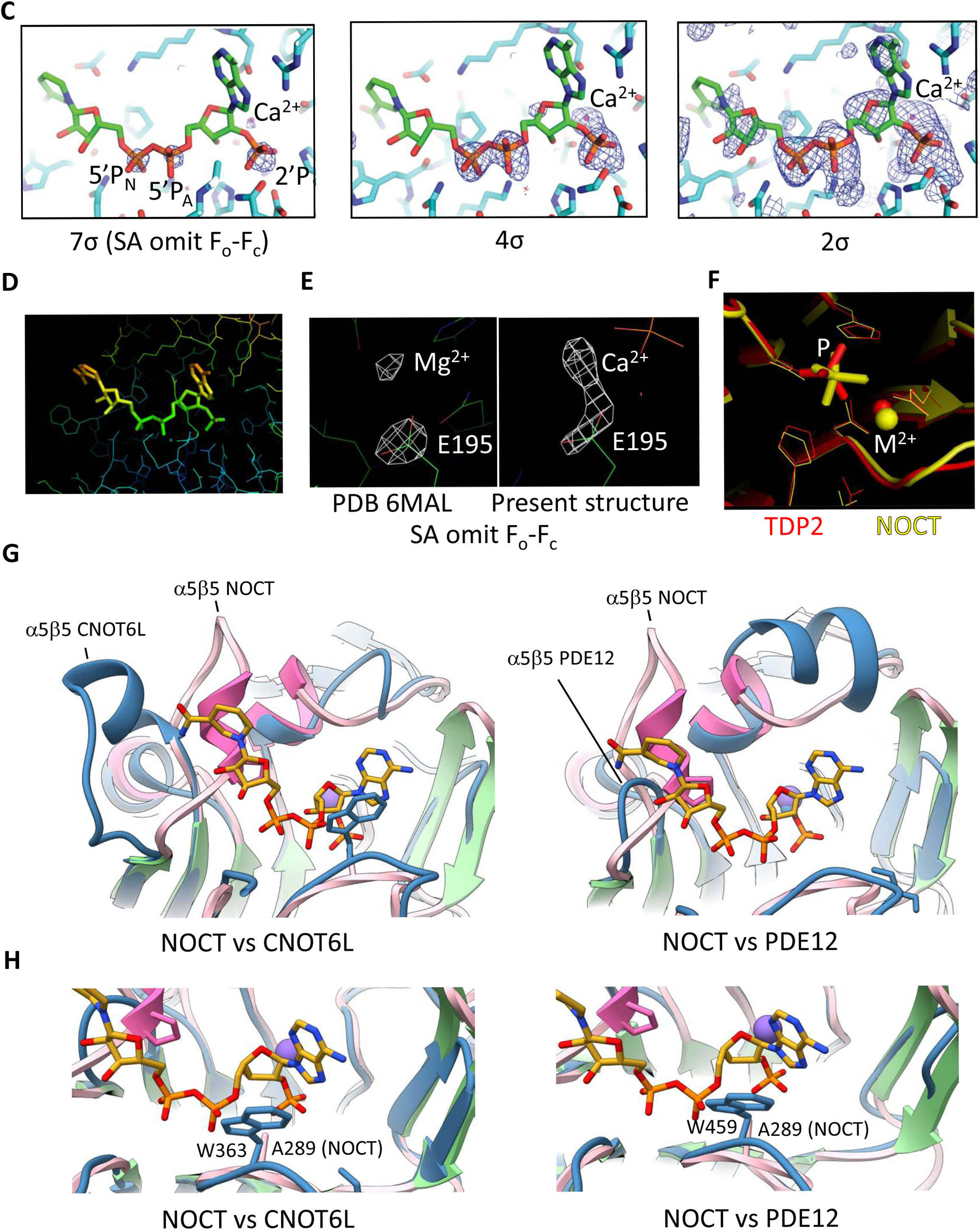
Diffraction properties of NOCT•NADPH crystals and structure comparisons with deadenylases. **A)** NOCT 122-431 activity is impaired by the addition of 2 mM CaCl_2_. Reactions were conducted for 30 minutes as in Fig. 2B. **B**) Completeness and CC_1/2_ plots for anisotropic diffraction data processing by STARANISO (http://staraniso.globalphasing.org/cgi-bin/staraniso.cgi). **C**) Simulated annealing (2000K) omit map for NADPH at three contour levels indicated on the figure. **D)** B-factor-colored NADPH (see also Table S1). **E**) Simulated annealing omit map for the previously published NOCT**•**Mg^2+^ complex 6MAL (Mg^2+^ and CD+OE1+OE2 atoms of E195 were omitted) vs a similar map for the NOCT**•**NADPH**•**Ca^2+^ complex (Ca^2+^ and CD+OE1+OE2 atoms of E195 were omitted). The electron density for Ca^2+^ is stronger than that for Mg^2+^, as expected due to higher electron count in Ca^2+^. **F**) Superposition of the TDP2 DNA repair enzyme in complex with 5’-P DNA (PDB ID 5INL) vs the NOCT**•**NADPH**•**Ca^2+^ complex. **G**) NOCT superimposition with CNOT6L and PDE12 showing α5β5 motifs. **H**) NOCT superimposition with CNOT6L and PDE12 showing a tryptophan in place of the NOCT alanine 289.

**Figure S5.**
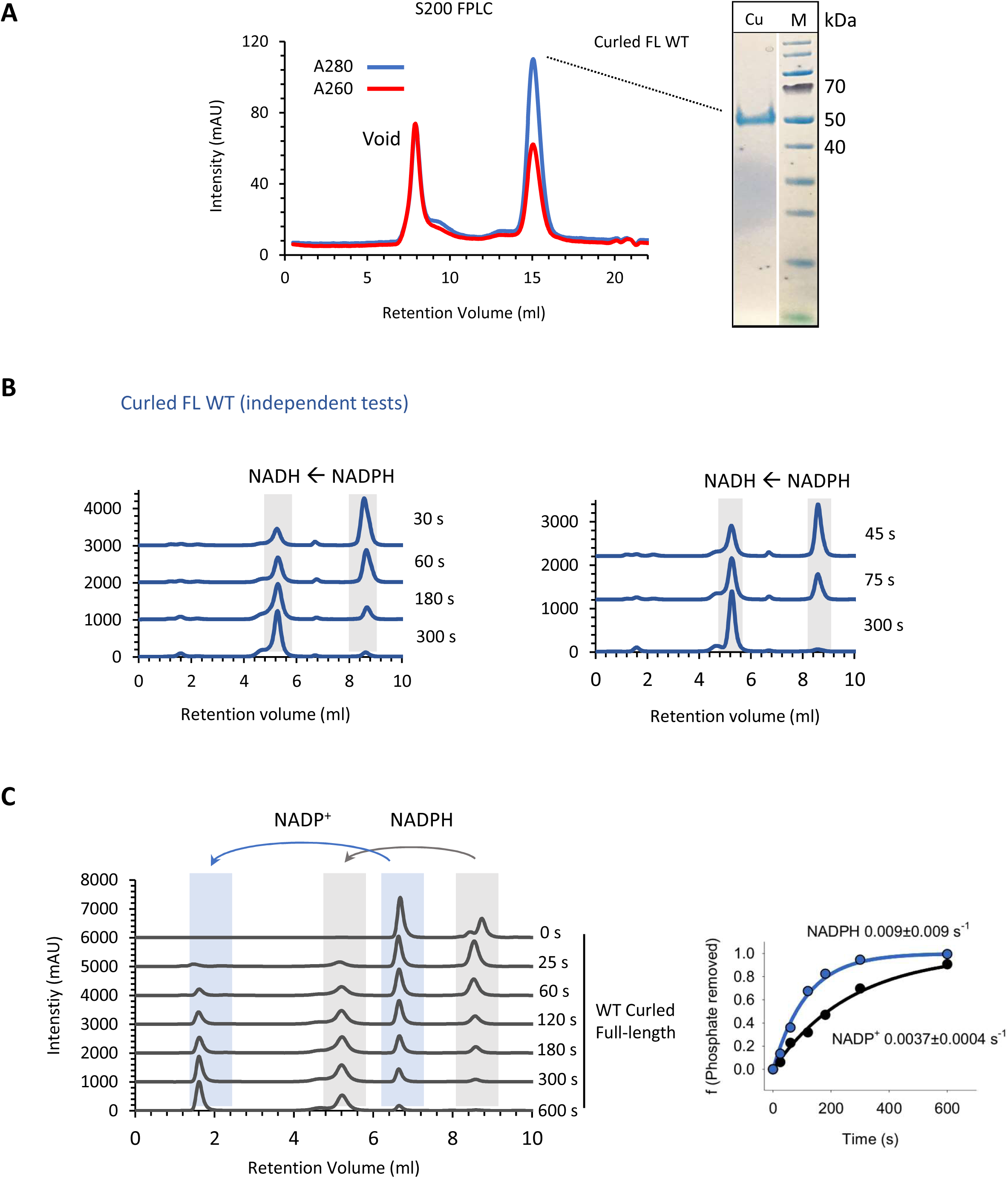
Curled purification and activity assay. **A**) S200 size exclusion purification trace for full length *Drosophila melanogaster* Curled protein and NuPAGE analysis of the peak. The protein was concentrated to 0.9 mg/ml (19.23 μM). **B**) Cleavage of NADPH (1 mM) by Curled (0.5 μM) related to Fig. 5C. **C**) Left: NADP^+^ and NADPH cleavage by Curled under conditions of kinetic competition as in Fig. S2A. Reactions contained 500 nM Curled and 1 mM of each nucleotide. Right: quantitation of the data fitted to single-exponential kinetics.

**Table S3.**
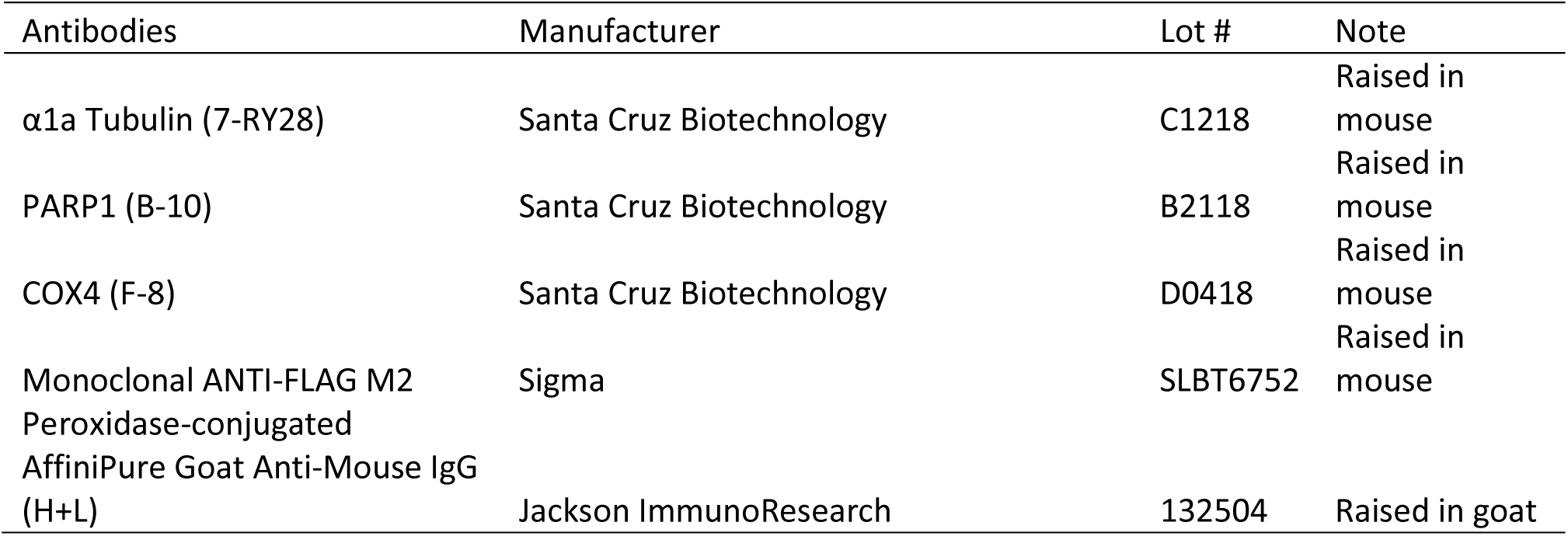
Antibodies used

**Table S2.**
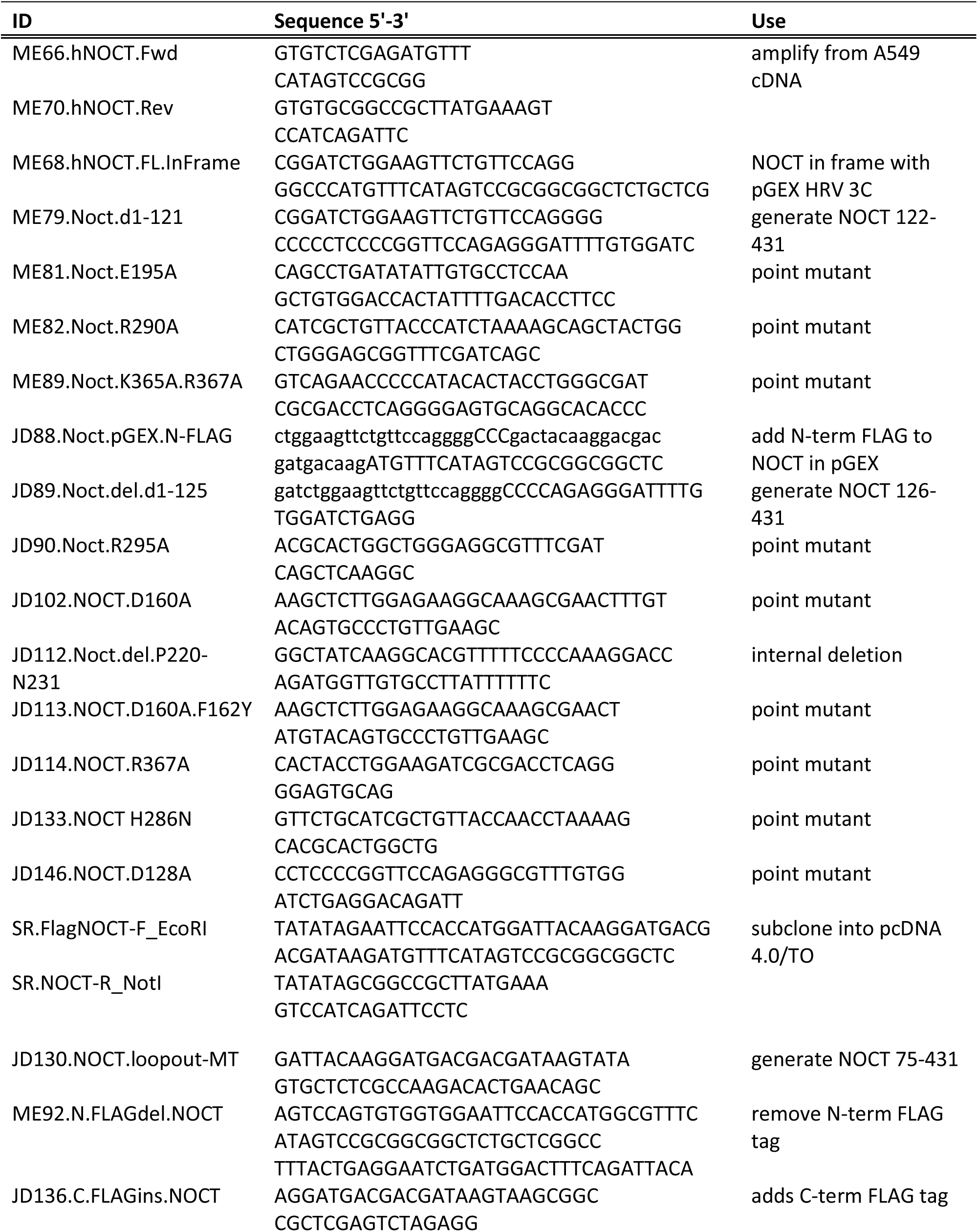
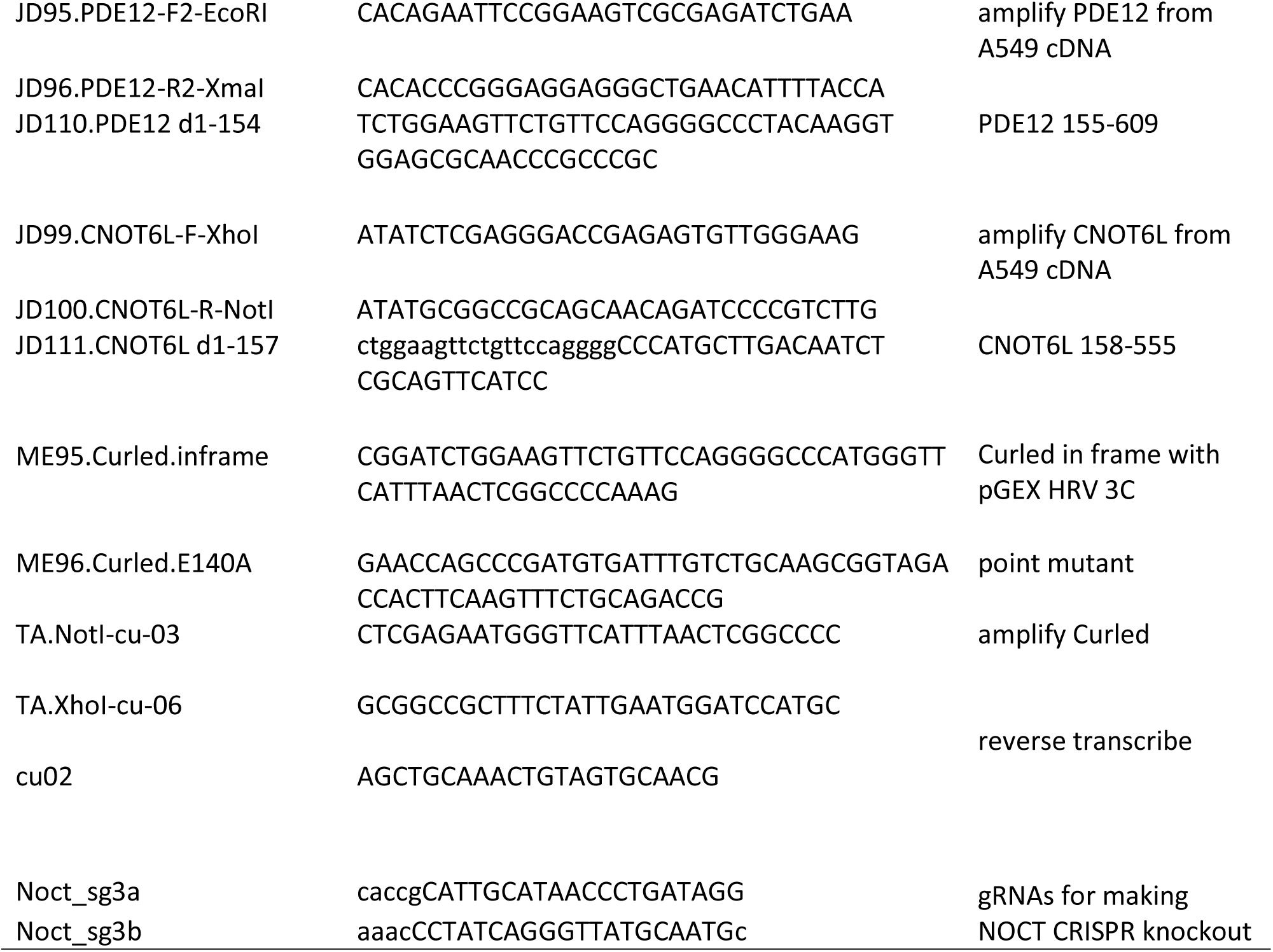
Oligonucleotide sequences.

**Table S1.** Data collection and refinement statistics

| Human Nocturnin 122-431 • NADPH • Ca<sup>2+</sup> |  |
| --- | --- |
| **Data collection** |  |
| Space Group | P4<sub>1</sub>2<sub>1</sub>2 |
| Cell dimensions |  |
| a, b, c (Å) | 62.88, 62.88, 153.59 |
| α, β, γ (°) | 90.0, 90.0, 90.0 |
| Resolution (Å) | 29.09-2.05 (2.33–2.05) |
| *R*<sub>pim</sub> | 0.034 (0.443) |
| CC(1/2) | 0.999 (0.760) |
| I/σI | 14.27 (1.86) |
| Completeness % | 93.20* (82.39)* |
| Redundancy | 12.65 (11.88) |
| **Refinement** |  |
| Resolution (Å) | 2.70 |
| Completeness % | 93.69* |
| No. unique reflections | 8,470 (523) |
| *R*<sub>work</sub>/*R*<sub>free</sub> | 0.2063/0.2630 |
| No. atoms | 2,429 |
| Protein | 2,362 |
| NADPH | 48 |
| Ca | 1 |
| Water | 18 |
| B-factors |  |
| Protein | 48.67 |
| NADPH | 73.86 |
| Ca | 50.79 |
| Water | 40.44 |
| R.m.s deviations |  |
| Bond lengths (Å) | 0.005 |
| Bond angles (°) | 0.802 |
| Ramachandran plot (%) |  |
| Favored | 97.28 |
| Allowed | 2.72 |
| Disallowed | 0.0 |

Highest-resolution shell is shown in parentheses.
\*Ellipsoidal completeness.

## Supplementary Methods

### Cloning

The coding region of full-length human Nocturnin (residues 1-431) was amplified by PCR from an in-house cDNA library using poly I:C transfected A549 human lung epithelial cells (obtained originally from Prof. Susan Weiss, UPenn propagated in our group as previously described, Chitrakar A. et al., and Korennykh A., PNAS 2019) to upregulate NOCT expression. The *Drosophila melanogaster* Curled cDNA (splicing form H) was cloned by RT-PCR from the RNA pool of wild-type fly embryos. Seven micrograms of total RNA from Oregon-R embryo was reverse-transcribed by using the primer cu-02 (Table S2) followed by the PCR reaction with forward (XhoI-cu-03) and reverse (NotI-cu-06) primers. The single band of PCR product was purified from agarose gel and then subcloned into pCRII vector (Invitrogen). The cDNA clone did not include any base changes that cause amino-acid alteration from Curled protein sequences in the FlyBase (https://flybase.org). The PCR products were cloned into pGEX-6P bacterial expression vector (GE Healthcare Life Sciences). Deletion and point mutants were generated by one-primer site-directed mutagenesis as previously described^1^. The constructs for human PDE12 155-609 and CNOT6L 158-555 were also previously described^1^. For mammalian expression, full-length NOCT, ΔMTS-NOCT (75-431), and full-length NOCT-E195A were subcloned into pcDNA^TM^4/TO vector (Life Technologies). All constructs used in this study were verified by DNA sequencing.

### Protein purification

pGEX-6P vectors encoding recombinant proteins were transformed into *Escherichia coli* BL21 (DE3)-CodonPlus RIPL (Agilent Technologies) and grown to an OD600 of 0.4 in Luria-Bertani medium at 37 °C followed by induction with 0.2 mM isopropyl-β-D-thiogalactopyranoside (IPTG) and overnight expression at 22 °C. The cells were pelleted at 4,600 × *g* for 20 min, resuspended in lysis buffer (20 mM HEPES pH 7.4, 1 M KCl, 2 mM MgCl_2_, 1 mM EDTA, 10% (vol/vol) glycerol, 5 mM DTT, and Roche cOmplete protease inhibitor), and lysed on EmulsiFlex C3 (Avestin).

Crude lysates were clarified by centrifugation at 35,000 × *g* for 30 min, at 4 °C. Clarified lysates were affinity-purified using glutathione Sepharose (GE Healthcare Life Sciences). The GST tag was removed with Prescission protease (GE Healthcare Life Sciences). NOCT 122-431, PDE12 155-609, CNOT6L 158-555 and Curled 1-419 variants were similarly purified. Full-length NOCT was further purified by MonoQ, then by MonoS ion-exchange chromatography, and lastly, by Superdex S200 size-exclusion chromatography. NOCT constructs were purified using MonoS, MonoQ and Superdex 200 size-exclusion chromatography. For PDE12, CNOT6L and Curled only size-exclusion chromatography was used. Size exclusion buffer contained 20 mM HEPES (pH7.4), 350 mM KCl, 1mM EDTA, 10% (vol/vol) glycerol, and 5mM DTT. All proteins were purified to ≥ 95% purity. Concentrations were determined by UV 280 spectrophotometry, using protein sequence for calculating extinction coefficients.

### Immuno-precipitation of human FLAG-NOCT from A549 cells

Human A549 cells grown in 10 cm dishes were transfected with 8 *µ*g pcDNA^TM^4/TO plasmids expressing either full length WT or E195A human NOCT, with a C-terminal FLAG tag. Transfections were carried out using Lipofectamine 2000 (Invitrogen) per manufacturer’s instructions, for 18 hours. Cells were washed once, scraped in cold PBS, transferred to 1.5 ml Eppendorf tubes and pelleted at 500 x g for 5 min (at 4 °C). The cell pellets were resuspended in lysis buffer (10 mM HEPES pH 7.5, 10 mM NaCl, 2 mM EDTA and 0.5% Triton X-100), supplemented with complete protease inhibitor cocktail (Roche), RNase inhibitor, then constantly rotated at 4 °C for 10 min. Cell lysates were centrifuged at 1000 x g for 10 min at 4 °C to pellet nuclei. Supernatants were incubated with anti-FLAG M2 magnetic meads (Sigma; the list of antibodies used in the work is provided in Table S3), which were pre-equilibrated with lysis buffer. After incubating beads with cell lysates at 4 °C for two hours, beads were washed three times for 2 min in ice-cold wash buffer (10 mM HEPES pH 7.5, 150 mM NaCl, 0.5% Triton X-100). Lastly, beads were washed and resuspended in reaction buffer (20 mM Tris•HCl pH 8.0, 70 mM NaCl and 2 mM MgCl_2_), and immediately used for assays. Beads incubated with naïve A549 cells lysates served as a control for non-specific interactions.

### Microscopy

A549 cells were seeded on a chamber slide (Lab-Tek) in RPMI-1640 medium (Sigma) supplemented with 10% FBS at 37 °C. Cells were transfected using Lipofectamine 2000 (Invitrogen) with pcDNA^TM^4/TO plasmids (Life Technologies) encoding either WT full length human NOCT, E195A full length human NOCT or ΔMTS human NOCT. Twenty four hours after transfection, cells were briefly washed with serum-free RPMI-1640 and stained with 5 μM MitoSOX™ Red (Invitrogen, Lot # 1985406) at 37 °C for 10 min. Cells were washed again with PBS, fixed with 4% paraformaldehyde in PBS at room temperature (15 min), permeabilized with 0.1% TX-100 in PBS (10 min), then blocked with 20% goat serum in PBS (20 min). Mouse anti-FLAG (1:500, M2 Sigma) was used as a primary antibody and AlexaFluor 488 goat anti-mouse (1:500, Life Technologies, Lot #1613346) was used as a secondary antibody. Cells were further stained with DAPI (1:1000, Thermo Scientific, Lot# PJ1919631) and images were taken with Nikon Instruments A1 Confocal Laser Microscope. Nikon Elements software was used to prepare the images.

### Subcellular fractionations and western blots

The fractionation protocol was adapted from Jean Beltran et al.^2^. Three 15 cm dishes with A549 cells were transfected using Lipofectamine 2000 (Invitrogen) with plasmids expressing human NOCT (WT, E195A or ΔMTS). Cells were harvested 24 hrs post transfection pelleted at 4 °C and re-suspended in lysis buffer (10 mM HEPES (pH 7.4), 10 mM NaCl, 2.5 mM MgCl_2_). The lysates were supplemented with cOmplete protease inhibitor cocktail (Roche) and placed on a rotator for 10 min. Resuspended cells were further disrupted with 20 strokes on a dunce homogenizer. Lysate were centrifuged at 700 x g for 10 min to pellet nuclei, which were kept for further processing. The supernatant was centrifuged at 20,000 x g for 30 min to pellet organelles, including mitochondria. The resulting supernatant, containing the cytosolic fraction, was kept for further processing.

The organelle pellet was resuspended in 0.25 M sucrose, 6 mM EDTA, 120 mM HEPES pH 7.4, and top-loaded in a discontinuous density gradient established by iodixanol (Sigma OptiPrep^TM^) at 0%, 10%, 15%, 20%, 25%, and 30%. The tubes were centrifuged at 130,000 x g for three hours using an SW 60-Ti rotor (Beckman). Fractions from the 15% and 20% layers were diluted with PBS then centrifuged at 20,000 x g for 30 min to pellet the mitochondria. The nuclear pellet and the mitochondrial pellet were resuspended in Tris-buffered saline and lysed by sonication. Total protein concentration in each fraction was determined by Bradford assay. Equal OD_595_ units were loaded and separated by 10% BisTris PAGE (NuPAGE). The gels were transferred to PVDF membranes (Life Technologies), followed by 15 minutes of blocking with 5% milk. The membranes were incubated with 1:1000 mouse anti-human PARP-1 antibody (Santa Cruz Biotech), 1:1000 mouse anti-human COX4 antibody (Santa Cruz Biotech), 1:1000 mouse anti-human α1a Tubulin antibody (Santa Cruz Biotech), or 1:1000 mouse anti FLAG (Sigma) primary antibody at 4 °C, overnight. The membranes were washed with Tris-buffered saline-Tween buffer and incubated with horseradish peroxidase-conjugated with anti-mouse secondary antibody (1:10,000, Jackson ImmunoResearch) for 30 minutes. The membranes were washed again and visualized using X-ray film and enhanced chemiluminescence western blotting detection reagents (GE Healthcare Life Sciences).

### Construction of A459 NOCT-KO cells

The oligonucleotide sequences used for the generation of small guide RNAs (sgRNA) are shown in Table S2. A pair of forward and reverse oligonucleotides for the generation of each sgRNA (synthesized by IDT) was annealed and inserted into plasmid vectors pLenti-CRISPRv2 (a gift Prof. Alex Ploss, Princeton University), between BsmBI restriction sites. For lentivirus production, HEK293T cells were plated into six-well plates to achieve 50-60% confluency in 24 hrs. On the next day, cells were transfected with 1.5 μg pLenti-CRIPSRv2 (with sgRNA), 1.33 μg pCMVdR8.91, and 0.17 μg of pMD2.G (a gift from Prof. Jared Toettcher, Princeton University), using FuGENE HD Transfection reagent (Promega) (9 μL in 150 μL of OptiMEM per well). Lentivirus-containing medium was collected after 48 hours. Following collection, the medium was passed through 0.45 μm filter. Polybrene 5 μg/mL (f/c) and HEPES (pH7.5; 100 mM f/c) were then added.

A549 cells at 40% confluency were infected with 100 μl of lentivirus-containing media in six well plates. The media was changed 24 hours post-infection, and puromycin (2 μg/ml f/c) was added three days post-infection. After four days of selection, surviving cells were cloned by limited dilution. Single-cell clones were selected for further amplification and genotyping and, independently, transcript-level mutation confirmation.

### P-value and error calculations

Statistical significance (P) was calculated using Welch’s two-tailed unpaired t-test. Error bars show S.E. as defined in figure legends.

### Radiolabeled RNA cleavage

RNA labeling with ^32^P and cleavage reactions were conducted as described previously^1^.

### RNA-seq

RNA-seq was conducted and analyzed as previously described^3^.

### GSEA analysis

GSEA analysis was conducted using GSEA data sets provided by the Broad Institute and generated from GO annotations as previously described^4^.

### X-ray structure visualization and analysis

Structure images were generated using PyMol (DeLano Scientific built) and UCSF Chimera^5^. Sequence conservation and 3D conservation mapping was conducted using SEQMOL-Kd (BiochemLabSolutions.com).

## References

1. Reppert, S. M. & Weaver, D. R. Coordination of circadian timing in mammals. Nature 418, 935–41 (2002).

2. Baggs, J. E. & Green, C. B. Functional analysis of nocturnin: a circadian clock-regulated gene identified by differential display. Methods Mol Biol 317, 243–54 (2006).

3. Baggs, J. E. & Green, C. B. Nocturnin, a deadenylase in Xenopus laevis retina: a mechanism for posttranscriptional control of circadian-related mRNA. Curr Biol 13, 189–98 (2003).

4. Gronke, S., Bickmeyer, I., Wunderlich, R., Jackle, H. & Kuhnlein, R. P. Curled encodes the Drosophila homolog of the vertebrate circadian deadenylase Nocturnin. Genetics 183, 219–32 (2009).

5. Green, C. B. et al. Loss of Nocturnin, a circadian deadenylase, confers resistance to hepatic steatosis and diet-induced obesity. Proc Natl Acad Sci U S A 104, 9888–93 (2007).

6. Estrella, M. A., Du, J. & Korennykh, A. Crystal Structure of Human Nocturnin Catalytic Domain. Sci Rep 8, 16294 (2018).

7. Wang, Y. et al. Rhythmic expression of Nocturnin mRNA in multiple tissues of the mouse. BMC Dev Biol 1, 9 (2001).

8. Niu, S. et al. The circadian deadenylase Nocturnin is necessary for stabilization of the iNOS mRNA in mice. PLoS One 6, e26954 (2011).

9. Stubblefield, J. J., Terrien, J. & Green, C. B. Nocturnin: at the crossroads of clocks and metabolism. Trends Endocrinol Metab 23, 326–33 (2012).

10. Stubblefield, J. J. et al. Temporal Control of Metabolic Amplitude by Nocturnin. Cell Rep 22, 1225–1235 (2018).

11. Pearce, S. F. et al. Maturation of selected human mitochondrial tRNAs requires deadenylation. Elife 6 (2017).

12. Wang, H. et al. Crystal structure of the human CNOT6L nuclease domain reveals strict poly(A) substrate specificity. EMBO J 29, 2566–76 (2010).

13. Abshire, E. T. et al. The structure of human Nocturnin reveals a conserved ribonuclease domain that represses target transcript translation and abundance in cells. Nucleic Acids Res 46, 6257–6270 (2018).

14. Chien-Kuang & Cornelia Ding, J. R., Jianli Wu, Tianai Sun, Kai-Yuan Chen, Po-Han Chen, Emily Xu, Sarah Tian, Jadesola Akinwuntan, Ziqiang Guan, Pei Zhou, Jen-Tsan Ashley Chi. Mammalian stringent-like response mediated by the cytosolic NADPH phosphatase MESH1. BioRxiv https://doi.org/10.1101/325266 (2018).

15. Ohashi, K., Kawai, S. & Murata, K. Identification and characterization of a human mitochondrial NAD kinase. Nat Commun 3, 1248 (2012).

16. Fukasawa, Y. et al. MitoFates: improved prediction of mitochondrial targeting sequences and their cleavage sites. Mol Cell Proteomics 14, 1113–26 (2015).

17. Kawai, M. et al. A circadian-regulated gene, Nocturnin, promotes adipogenesis by stimulating PPAR-gamma nuclear translocation. Proc Natl Acad Sci U S A 107, 10508–13 (2010).

18. Calvo, S. E., Clauser, K. R. & Mootha, V. K. MitoCarta2.0: an updated inventory of mammalian mitochondrial proteins. Nucleic Acids Res 44, D1251–7 (2016).

19. Subramanian, A. et al. Gene set enrichment analysis: a knowledge-based approach for interpreting genome-wide expression profiles. Proc Natl Acad Sci U S A 102, 15545–50 (2005).

20. Jiao, X. et al. 5’ End Nicotinamide Adenine Dinucleotide Cap in Human Cells Promotes RNA Decay through DXO-Mediated deNADding. Cell 168, 1015–1027 e10 (2017).

21. Wood, E. R. et al. The Role of Phosphodiesterase 12 (PDE12) as a Negative Regulator of the Innate Immune Response and the Discovery of Antiviral Inhibitors. J Biol Chem (2015).

22. Canto, C., Menzies, K. J. & Auwerx, J. NAD(+) Metabolism and the Control of Energy Homeostasis: A Balancing Act between Mitochondria and the Nucleus. Cell Metab 22, 31–53 (2015).

23. Smeland, T. E., Nada, M., Cuebas, D. & Schulz, H. NADPH-dependent beta-oxidation of unsaturated fatty acids with double bonds extending from odd-numbered carbon atoms. Proc Natl Acad Sci U S A 89, 6673–7 (1992).

24. Li, Y. et al. Deficiency in the NADPH oxidase 4 predisposes towards diet-induced obesity. Int J Obes (Lond) 36, 1503–13 (2012).

25. Jiang, F. et al. Systemic upregulation of NADPH oxidase in diet-induced obesity in rats. Redox Rep 16, 223–9 (2011).

26. Carroll, M. F. & Schade, D. S. The dawn phenomenon revisited: implications for diabetes therapy. Endocr Pract 11, 55–64 (2005).

27. Hurd, T. R., Liang, F. X. & Lehmann, R. Curly Encodes Dual Oxidase, Which Acts with Heme Peroxidase Curly Su to Shape the Adult Drosophila Wing. PLoS Genet 11, e1005625 (2015).

28. Gasteiger, E. et al. ExPASy: The proteomics server for in-depth protein knowledge and analysis. Nucleic Acids Res 31, 3784–8 (2003).

## References

1. Estrella, M. A., Du, J. & Korennykh, A. Crystal Structure of Human Nocturnin Catalytic Domain. Sci Rep 8, 16294 (2018).

2. Jean Beltran, P. M., Mathias, R. A. & Cristea, I. M. A Portrait of the Human Organelle Proteome In Space and Time during Cytomegalovirus Infection. Cell Syst 3, 361–373 e6 (2016).

3. Alisha Chitrakar et al. Realtime 2-5A kinetics suggests interferons β and lambda evade global arrest of translation by RNase L. BioRxiv https://doi.org/10.1101/476341 (2018).

4. Rath, S. et al. Human RNase L tunes gene expression by selectively destabilizing the microRNA-regulated transcriptome. Proc Natl Acad Sci U S A 112, 15916–21 (2015).

5. Pettersen, E. F. et al. UCSF Chimera--a visualization system for exploratory research and analysis. J Comput Chem 25, 1605–12 (2004).

